# Application of the fluctuation theorem for non-invasive force measurement in living neuronal axons

**DOI:** 10.1101/233064

**Authors:** Kumiko Hayashi, Yuta Tsuchizawa, Mitsuhiro Iwaki, Yasushi Okada

**Affiliations:** Department of Applied Physics, Graduate School of Engineering, Tohoku University, Sendai, Japan; Laboratory for Cell Polarity Regulation, Center for Biosystems Dynamics Research (BDR), RIKEN, Osaka, Japan; Graduate School of Frontier Biosciences, Osaka University, Osaka, Japan; Laboratory for Cell Dynamics Observation, Center for Biosystems Dynamics Research (BDR), RIKEN, Osaka, Japan; Department of Physics, Universal Biology Institute (UBI), and the International Research Center for Neurointelligence (WPI-IRCN), The University of Tokyo, Tokyo, Japan

## Abstract

Although its importance is recently widely accepted, force measurement has been difficult in living biological systems, mainly due to the lack of the versatile non-invasive force measurement methods. The fluctuation theorem, which represents the thermodynamic properties of small fluctuating non-equilibrium systems, has been applied to the analysis of the thermodynamic properties of motor proteins *in vitro*. Here, we extend it to the axonal transport (displacement) of endosomes. The distribution of the displacement fluctuation had three or four distinct peaks around multiples of a unit value, which the fluctuation theorem can convert into the drag force exerted on the endosomes. The results demonstrated that a single cargo vesicle is conveyed by one to three or four units of force production.

## Introduction

One of the technical hurdles in mechanobiology, a growing field of science at the interface of biology and physics, has been the methods to measure force in living cells non-invasively. The force or stress on the outer surface of the cells, or the plasma membrane, can be measured by traction force microscopy (Polacheck and Chen, 2016). Fluorescent protein-based biosensors for force or tension at the cellular levels are also actively being developed using Förster resonance energy transfer (Meng *et al*., 2008; Meng and Sachs, 2012; Guo *et al*., 2014). Optical tweezers have been used to measure force exerted on the lipid droplets in cultured cells or in Drosophila embryos (Shubeita *et al*., 2008; Jun *et al*., 2014; Mas *et al*., 2014), but their application to other organelles or subcellular structures is difficult. Stokes’ law can be theoretically used to estimate the drag force on the organelles moving at the velocity *v* as *F*=*6πηrv*. But, the viscosity *η* and the diameter of the organelle *r* need to be measured, the latter of which is often difficult for small organelles whose size is close to or below the diffraction limit of the microscope resolution.

A good example that needs force measurements *in vivo* is the axonal transport of vesicles, which are transported mainly by kinesins from the cell body to the periphery (anterograde) and dynein for the reverse direction (retrograde) (Hirokawa *et al*., 2009). Although many studies to date have elucidated their biological or functional importance, many physical or biophysical properties are still unclear (Encalada and Goldstein, 2014). For example, there is still controversy regarding the relationship between motor number, velocity and force (Figure 1A). The *in vitro* studies of purified kinesin-1 have established that its velocity is around 1 μm/s without external load force. The velocity does not decrease much under low load forces (up to around 2 pN) and decreases with higher load force to zero at around 10 pN, which is called the stall force. The exact value for the stall force of kinesin ranges from 5 to 10 pN depending on the kinesin isoforms and experimental conditions. In this paper, however, we assume the stall force of kinesin as 10 pN for the analysis of the force values. Thus, kinesin can carry a micro-meter sized bead at around 1 μm/s in vitro, because the drag force on the bead in water is much smaller than 1 pN (blue line in Figure 1A). Contrastingly, cytoplasm is densely filled with proteins and its viscosity is 1000 times larger than water (Wirtz, 2009; Hayashi *et al*., 2013). Thus, the drag force on the axonally transported vesicle would be about a few pico-newtons or more, and its velocity would be slowed down (purple line in Figure 1A). However, the cargo velocity in the axon is often much faster than the maximum velocity in vitro and is as fast as 5 μm/s (Allen *et al*., 1982; Chiba *et al*., 2014). Mammalian dynein is even more controversial. Many studies report its maximum force as only around 1 pN (Mallik *et al*., 2004; Hendricks *et al*., 2012; Rai *et al*., 2013), though a few studies report values (5-10 pN) close to that of kinesin (~10 pN) (Toba *et al*., 2006; Nicholas *et al*., 2015; Belyy *et al*., 2016).

**Figure 1.**
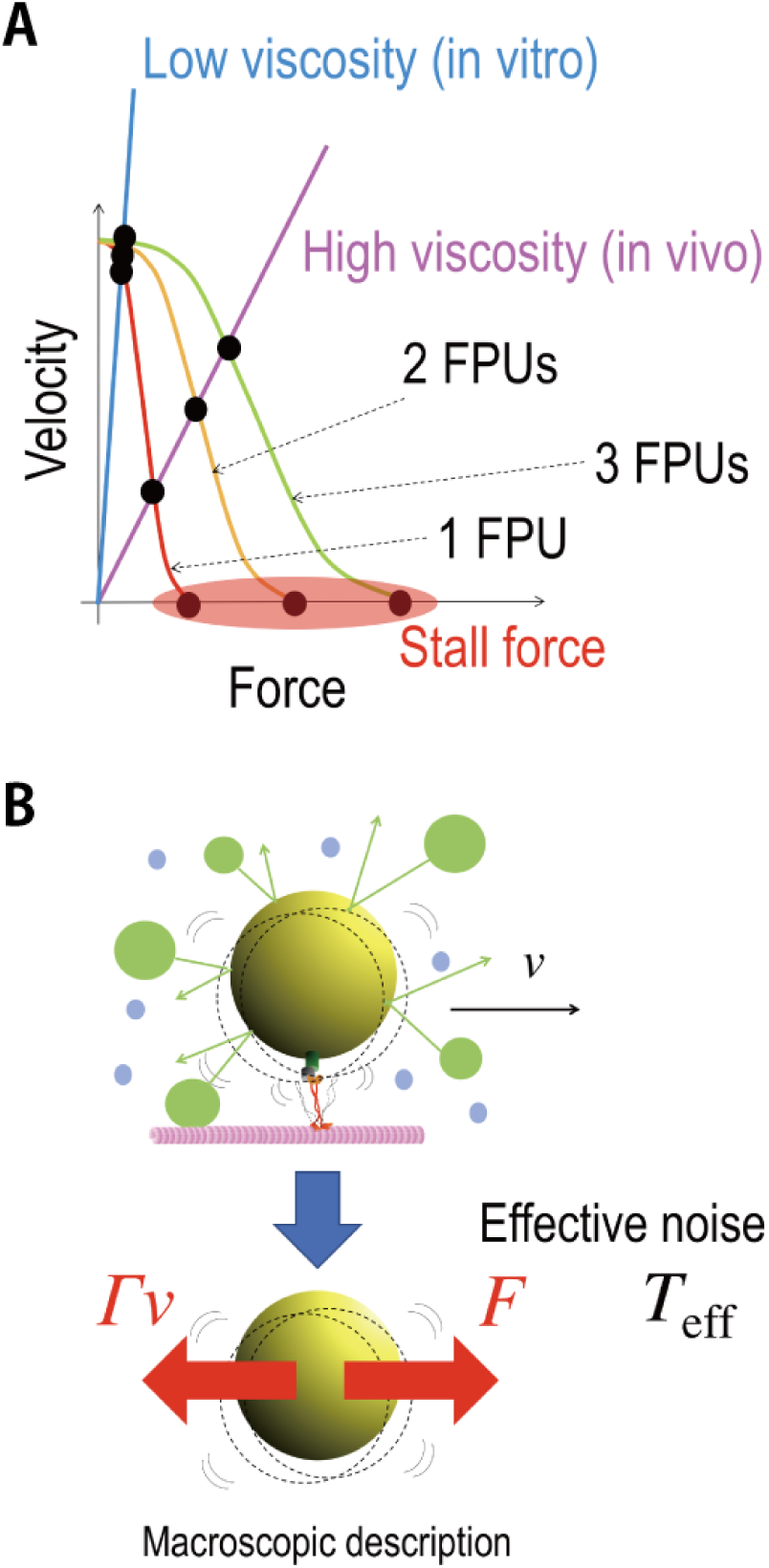
Schematics of the theoretical idea of FT. **(A)** The schematics of the force-velocity relations for motor proteins (=force producing units (FPUs)). The curves represent the case of 1FPU (red), 2FPUs (yellow) and 3FPUs (green). The lines represent the relation *Γv*=*F* where *F* is a force exerted by motors, *v* and *Γ* are the velocity and friction coefficient of a cargo vesicle. The values of drag force for 1FPU, 2FPUs and 3FPUs are considered to be discrete in the high viscous case (purple line) similar to the values of stall force. **(B)** A cargo (yellow) is transported by motors (red) along a microtubule (pink) at a constant velocity (*v*) (top). Here the dashed circles represent fluctuating movement of the cargo, and the blue and green circles represent water molecules, vesicles or organelles that collide with the cargo. The cargo fluctuates due to the collision with other vesicles and organelles in addition to thermal noise, while exhibiting directional motion. In the macroscopic description of the cargo transport (bottom), force (*F*) exerted by motors is balanced by the drag force (*Δv*). In the macroscopic description, the fluctuation of the cargo is described by the effective temperature (*T*_eff_) representing the additional noise. In this paper, *F* is estimated from this fluctuation via the FT.

To investigate these questions, we wanted to measure the drag force on axonally transported vesicles. There are several previous reports on the measurement of the force with vesicles or lipid droplets in living cells using optical tweezers. However, those experiments only measured the responses (slowing down or stalling) to the external load force applied by the optical tweezers. The stall force can be measured but it is not the drag force on the vesicle during the transport. To estimate the drag force, here we propose a method using the fluctuation theorem (FT). FT is a new universal law for entropy production in small non-equilibrium systems actively studied in the field of physics, which connects energy dissipation to fluctuation (Ciliberto *et al*., 2010). In previous studies, for example, we have established that the FT can be applied to estimate molecular energies from the fluctuation property of bio-molecules *in vitro* (Hayashi *et al*., 2010). This approach enables the estimation of the energy or force from only the passive measurement, the fluctuation of the movement. Thus, it is a fully passive and non-invasive method, potentially suitable for measurements in living cells.

## Results

### Theory

For a colloidal bead moving at a constant speed (*v*) *in vitro*, under highly damped conditions, the motion of center position *X*(*t*) can be described by the Langevin equation:

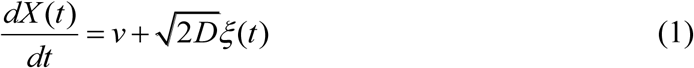

where the second term on the right-hand side represents the stochastic collisions with solvent molecules, where ξ is Gaussian noise with 〈*ξ*(*t*)*ξ*(*t*′)〉 = *δ*(*t* – *t*′), where < > denotes the time average over the time course, and *D* is a diffusion coefficient. It is well known, in near equilibrium, that *D* in equation (1) satisfies the fluctuation-dissipation theorem (Howard, 2001) *D*=*k*_B_*T*/*γ*, where *γ* is the friction coefficient of the bead, *k*_B_ is the Boltzmann constant, and *T* is the temperature of the environment. Using this relation, equation (1) is rewritten as

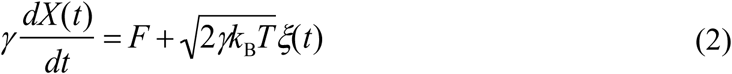

where *F* is a force defined as *F* =*γv*.

In the case of cargo vesicle transport in cells, these relations cannot be applied directly. There can be more interactions between the vesicle with the surrounding environments than the simple collisions with solvents. Vesicle transport might be hindered by the tethering interaction between the vesicle and the cytoskeleton, by the associating proteins on the microtubule or by the collision with other soluble proteins, cytoskeletons or organelles. Even in such complex situations, the friction force of the vesicle can be regarded to be proportional to the velocity unless the movement is too fast. This apparent frictional effect is characterized by the effective friction coefficient *Γ* that includes the interaction effects between the cargo and surrounding environment, so that it would be larger than *γ*=6*πηr* as assumed in the classical Stokes’ law (*r*: the radius of the cargo, *η*: the viscosity of the cytosol). This assumption was already examined experimentally by tracking the trajectories of the passive tracer beads in the cytoplasm (Wirtz, 2009).

Similarly, the second term on the right-hand side in equation (2) should be revised. Since the assumption for Gaussian noise implies that the noise is caused by large numbers of independent interactions, it is not dependent on whether the system is close to the thermal equilibrium or not. In fact, force noise acting on cargo vesicles measured in this study showed the statistical property of Gaussian noise judging from the probability density distributions and the power spectrum densities shown in the results sections below (Figure 2C, D). For the amplitude of the Gaussian noise, we replace the equilibrium temperature *T* with an empirical parameter, called effective temperature *T*_eff_ so as to include the additional noise caused by the complex interactions involved in cargo transport other than thermal noise (see the schematics in Figure 1B). Note that the idea of effective temperature was theoretically studied recently (Hayashi and Sasa, 2004; Cugliandolo, 2011) and was introduced to analyze the non-equilibrium motion of DNA molecules in a single-molecule experiment (Dieterich *et al*., 2015).

**Figure 2.**
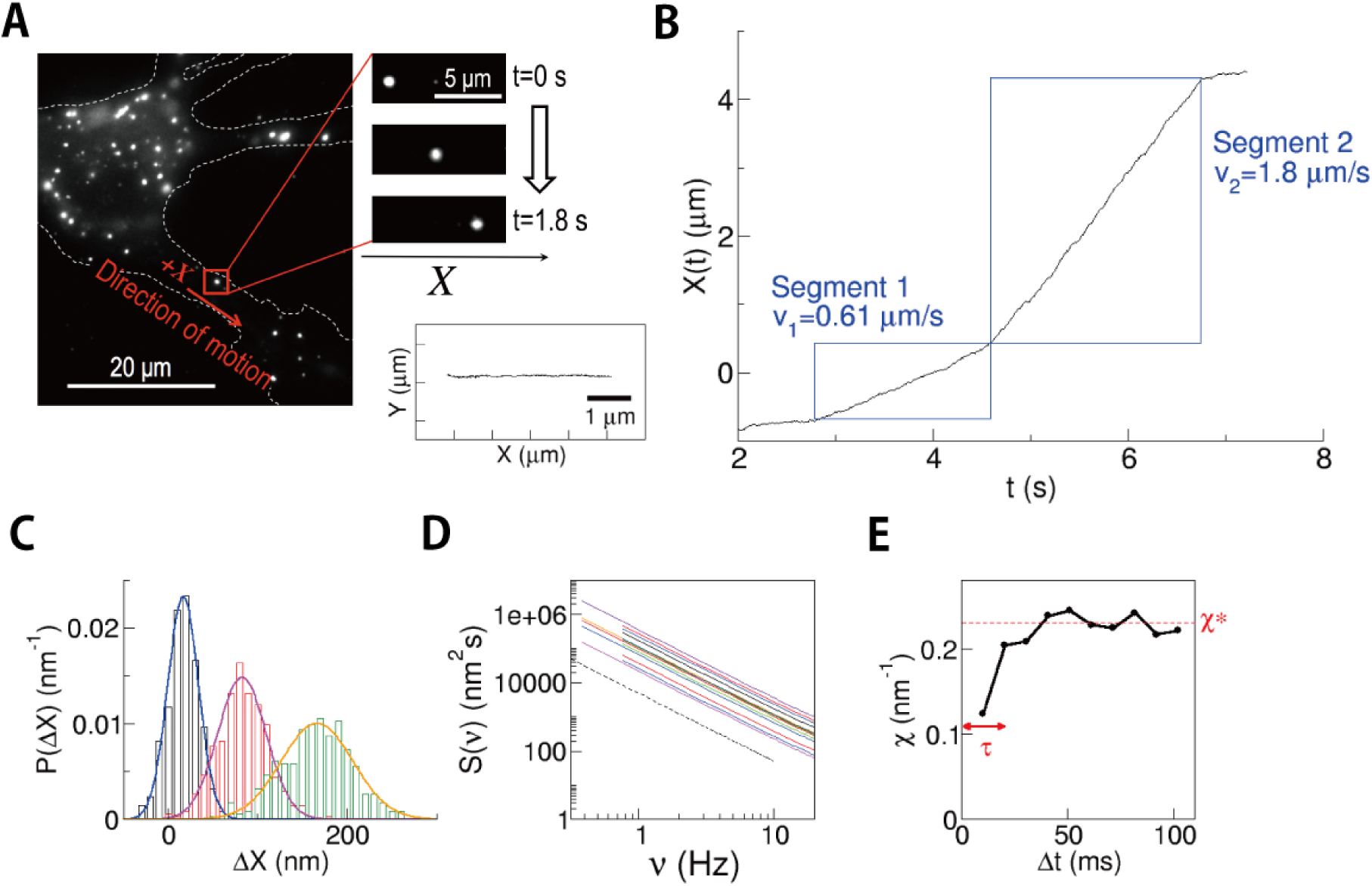
Measurement in the living neuron. **(A)** A typical view of DiI-stained endosomes in a SCG neuron. In the right bottom panel, a typical 2D-trajectory of an endosome during 4 s is shown. **(B)** An example of the trace of the displacement *X*(*t*) of an endosome, which showed a velocity change in the middle of the run event. The direction of movement was set as a plus direction of *X*. Segments for the analysis (rectangle areas) were determined by fitting the trajectory to the constant velocity movement as detailed in the Methods section. **(C)** Gaussian distribution of displacement *ΔX*=*X*(*t*+*Δt*)*-X*(*t*) in the cases *Δt*=10.2 ms (blue), 51.0 ms (pink), and 102 ms (green). (**D**) Power spectrum of the position *X*(*t*) in a constant velocity segment is inversely proportional to the square of the frequency (the dotted line with a slope of ‐2) (n=12), which is consistent with the assumption of Gaussian noise. **(E)** Relaxation of *χ. χ*\* and *τ* denote the value of convergence and the time constant, respectively. See also Supplemental Figure S1, S2 for the detailed analysis of *χ*.

With these modifications, the movement of a vesicle in the cytoplasm (Figure 1B) can be modeled as

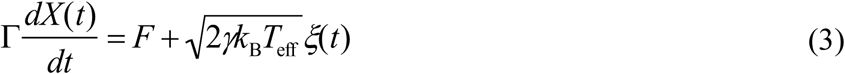

From the stochastic model (3), the following fluctuation theorem is derived (Hayashi *et al*., 2010):

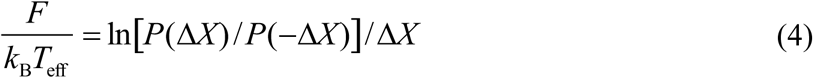

In this paper, the quantity on the right-hand side is called the degree of fluctuation *χ*:

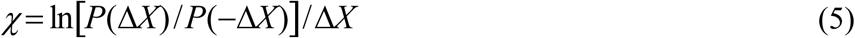

Then, equation (4) becomes

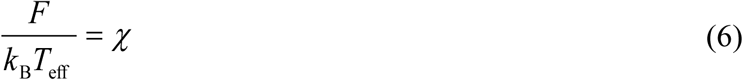

Here, it should be noted that equation (3) is a phenomenological macroscopic model. Microscopically, the motion of the vesicle, for example, will be described by a more complex model like

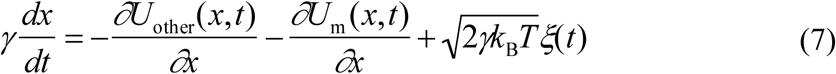

where 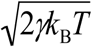 is thermal noise acting on the cargo vesicle, *U*_m_ is an interaction between vesicle and motors, *U*_other_ is the other interactions acting on the vesicle. Equation (3) corresponds to the coarse-grained model of equation (7), and the microscopic effect of *U*_m_ and *U*_other_ contributes to the effective viscosity *Γ*, effective noise 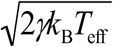 and the drag force *F* (Hayashi and Sasa, 2004, 2005).

Equation (6) holds for the time scale long enough to be compared with the characteristic time scales for the microscopic interactions *U*_m_ and *U*_other_. Hence, the right-hand side of equation (6), *χ* will show a relaxation behavior against the time interval *Δt* with the analysis for *ΔX*. The relaxation time constant *τ* is expected to show dependence on the cycle time of the motor protein, which was confirmed experimentally in the subsequent sections. Therefore, equation (6) is modified as follows to reflect that the converged value *χ*\*, instead of the transient values of *χ*, should be used for the estimation of the drag force:

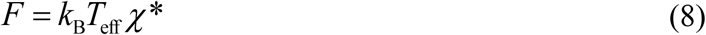

### Fluctuation measurement of axonally transported vesicles *in vivo*

We examined the FT (equation (8)) with a real biological system, the axonal transport of endosomes in supracervical ganglion (SCG) neurons. The endosomes were stained with a membrane-staining dye, DiI (Figure 2A). As established previously, most endosomes show linear movement along the axon anterogradely (to the axon terminal) or retrogradely (to the cell body). Although they sometimes show stochastic switching of the velocity or reversal of the direction, the fluctuation analyzed here is the fluctuation of the displacement around a constant velocity. Thus, the segment of unidirectional movement of constant velocity was chosen for the analysis (boxed regions in Figure 2B).

The movement of the endosome was recorded at the frame rate of 98 frames per sec. The position of the endosome was determined as the centroid of the fluorescent spot with the accuracy of 8 nm (see Methods for details). The degree of fluctuation *χ* (equation (5)) was calculated from the distribution *P*(*ΔX*) (Figure 2C) of the displacement during the time interval *Δt*, namely *ΔX*(*t*)= *ΔX*(*t*+*Δ*(*t*)- *ΔX*(*t*) as described above (see Methods for details, and Supplemental Figure S1 for the evaluation of the errors). Here we note that the assumption of Gaussian noise in *X*(*t*) assumed in the Theory section was verified by the Gaussian-like probability density distribution of *P*(*ΔX*) (Figure 2C) and the Brownian-noise power spectrum density *S*(*ν*) of the position *X*(*t*) (Figure 2D).

The degree of fluctuation *χ* thus calculated showed convergence (*χ*\*) but with a relaxation time (Figure 2E, Supplemental Figure S2A). The relaxation time did show dependency on the enzymatic turnover rate (Supplemental Figure S2B, C), but the relaxation time was much longer than the enzymatic cycle time (~10 ms/molecule with saturating ATP for kinesin). The microenvironment around the vesicle, especially its viscoelastic nature, would affect the relaxation time as explained in the Theory section.

### Velocity ratio

To validate the FT (equation (8)), we first searched for the traces that contain two successive constant velocity segments (~2 s duration for each segment) with different velocities as shown in Figure 2B, because such traces would enable us to test the FT without further estimation of the parameter values.

For each vesicle, its size or surrounding environment of the axon will not change substantially during the few seconds of the run event. Then, the friction coefficient *Γ* would be same for both before and after the velocity change. The drag forces in the two velocity segments are written as *F*_1_=*Γv*_1_ and *F*_2_=*Γv*_2_, respectively, given by Stokes’ law. If the FT (equation (8)) holds with the same *T*_eff_ value for both segments, *F*_1_=*k*_B_*T*_eff_ *χ*1* and *F*_2_=*k*_B_*T*_eff_ *χ*2*. Their ratio thus gives unity,

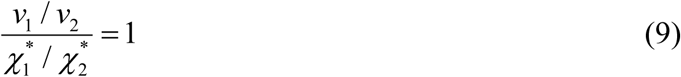

The result (Figure 3) was consistent with this relation, suggesting that the effective temperature would take the same or similar values for each vesicle during the few seconds of the run event. Here, it should be noted that the ratio 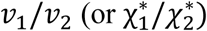 is equal to the ratio of the drag forces *F*_1_/*F*_2_ and it took values around 1/2, 2, 3, and 5, which suggests the discrete changes of the drag force as discussed below. This discrete behavior was previously reported for organelle transports in living cells (Levi *et al*., 2006; Shtridelman *et al*., 2008). Moreover, the proportional relation between *v* and *χ* was also examined with the kinesin-1 coated beads in vitro, where the optical tweezers were used to exert external load instead of viscous cytoplasm (Supplementary Figure S3).

**Figure 3.**
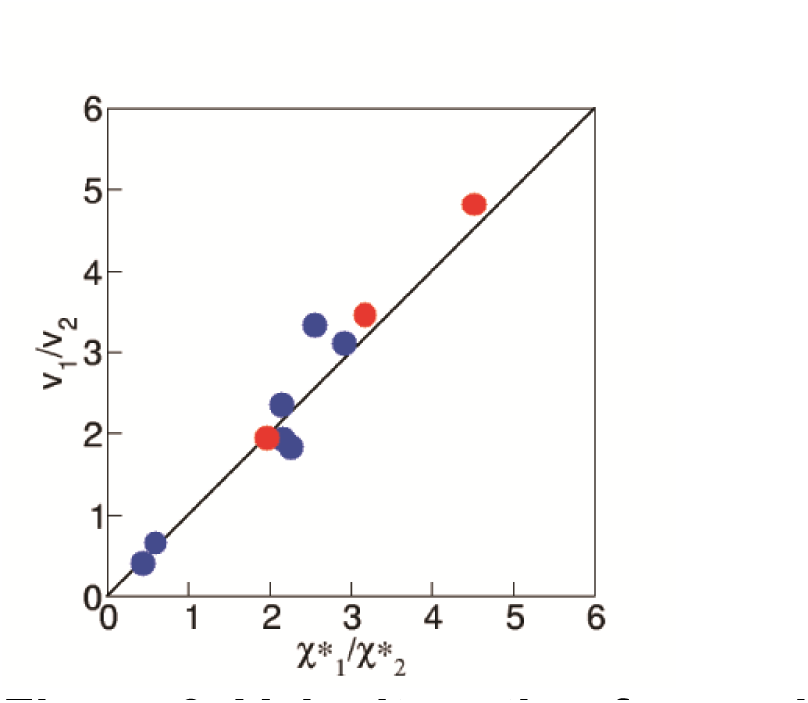
Velocity ratio of axonal endosomes *in vivo*. Confirmation of the proportional relation between the velocity and the fluctuation (equation (9)) (n=3 for anterograde, n=7 for retrograde from > 200 traces). The traces with two successive constant velocity segments with different velocities *v*_1_ and *v*_2_ (see Figure 2B for example) were analyzed.

### Fluctuation measurement for anterogradely transported endosomes in axons

Under the validation of equation (9), we then analyzed the remaining traces for the anterogradely transported endosomes with the segments of constant velocity that lasted for longer than 3 seconds. The measure of the fluctuation *χ* was calculated for the constant velocity region for each endosome for various intervals *Δt* from 10 ms to 100 ms, which confirmed that the time constant for the convergence was around *Δt*=50 ms (Figure 2E, 4A). We, therefore, analyzed the remaining shorter traces which had segments of constant velocity that lasted for about 2 seconds, with the interval *Δt* up to 50 ms.

**Figure 4.**
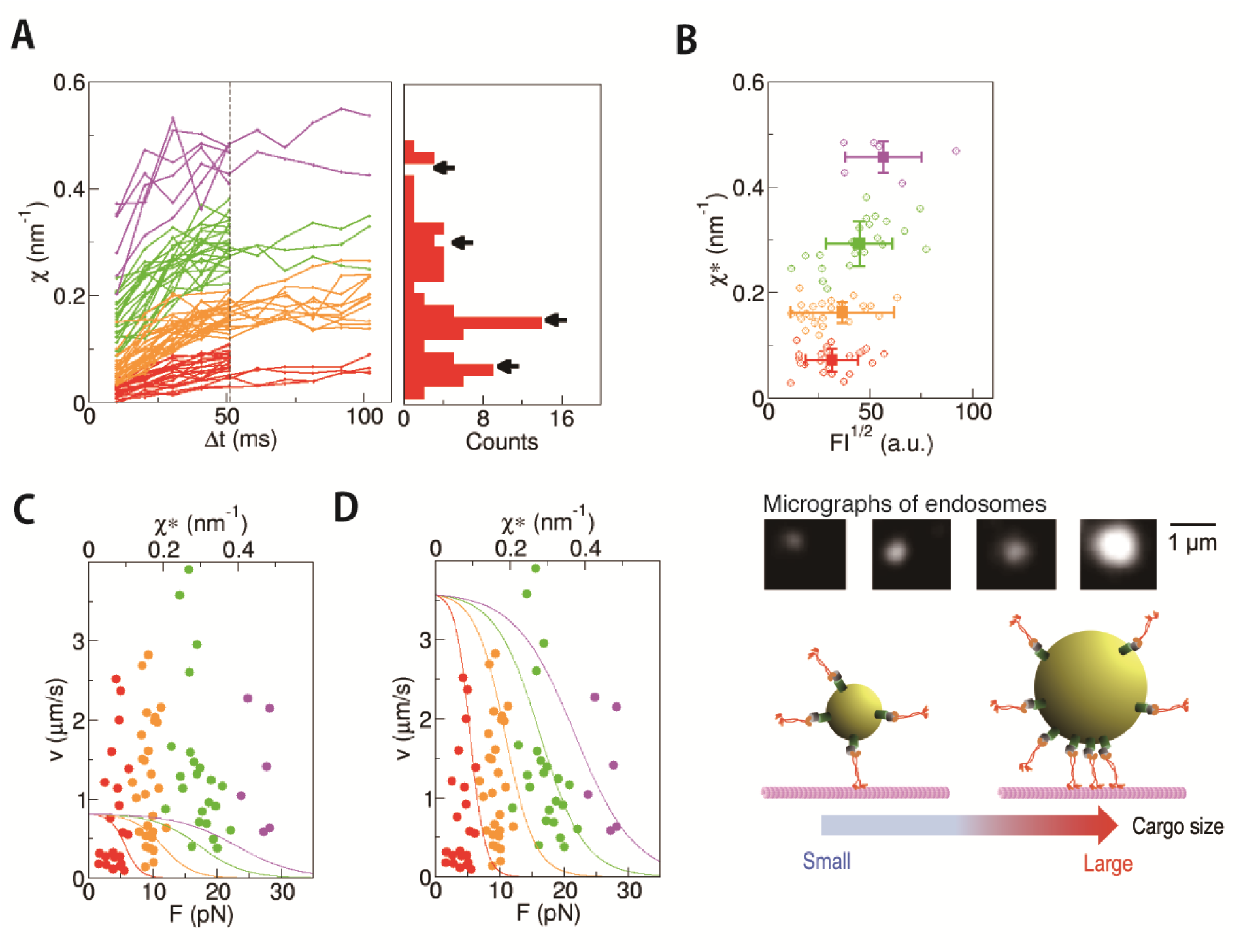
Measurement of χ for anterogradely transported endosomes in axon. **(A)** The traces of *χ* plotted against *Δt* for anterograde endosomes (n=79). They were classified into four clusters by *k*-means clustering (indicated with different colors. See Methods for the details). The distribution of *χ*\* values (*χ* at *Δt*=51 ms) is shown as a histogram. The positions of the cluster centers are indicated by arrows. **(B)** The fluctuation *χ*\* is plotted against the square root of fluorescence intensity (FI), which is a proxy for the radius of the endosome (middle panel). The color of each data point is the same as in the left panels in (A), which reflects the number of FPUs. The average and the SD for each cluster are shown with square symbols with error bars. There was a weak correlation between *χ*\* and (FI)^1/2^ (*r*=0.40), suggesting a tendency toward the larger endosomes experiencing a larger force with a greater number of force producing units (schema on the bottom). **(C, D)** For each endosome, the velocity is plotted against *χ*\*. The data points (circles) are plotted with the same colors as the panel (A). The curves are the results of *in vitro* measurements of kinesin-1 (Schnitzer *et al*., 2000) without adjustment of the maximum velocity (C) and after adjustment of the maximum velocity (D).

As summarized in Figure 4A, the plots (79 runs from 76 neurons) appeared to be clustered into several groups. The histogram of the *χ* values at *Δt*=50 ms, the proxy for the convergent value *χ*\*, showed discrete distribution, which was statistically confirmed by the *k*-means clustering (Methods). When *χ*\* is approximately proportional to force *F* (equation (8)), this discrete distribution of *χ*\* most likely reflects the force producing unit (FPU) in this system (see the schematics in Fig. 1A). Otherwise, the discrete force distribution by the presence of FPU would be obscured.

From the analysis of fluorescence intensities of the endosomes, a weak tendency toward more FPUs on larger endosomes was found (Figure 4B). The number of FPUs would thus not be tightly regulated to compensate for the greater drag resistance for the larger vesicles, but would simply reflect the geometric constraints that larger vesicles have more space to accommodate additional FPUs (Figure 4B, the bottom panel). It should be noted here that the velocity distribution does not show clear peaks (Supplemental Figure S4), probably reflecting the large variance of *Γ* due to the various sizes of the axonal endosomes (Figure 4B).

### Force-velocity relations for anterogradely transported endosomes in axons

For the anterogradely transported endosomes, the drag force should be balanced with the force produced by the motor protein, most likely kinesin (Figure 1B). Thus, the measured values for the force and the velocity should scatter along the force-velocity relation curve as determined by the mechanochemical properties of the motor protein kinesin.

For the anterogradely transported endosomes, one kinesin dimer molecule would most likely correspond to the anterograde FPU, because the previous biochemical measurement reported that only 1-4 kinesin dimers are bound to vesicles (Hendricks *et al*., 2010). Based on this assumption, we have compared the mechanical properties of kinesin *in vitro* and the force-velocity relation of the anterogradely transported endosome. Because the velocity was about 4 times faster *in vivo*, the *in vitro* force-velocity relation of kinesin (Schnitzer *et al*., 2000) did not match the result i*n vivo* at all (Figure 4C). Better fitting was achieved by increasing the enzymatic turnover rate (Figure 4D), which might suggest the acceleration of the enzymatic reaction due to the macromolecular crowding in the cytoplasm (Ellis, 2001) or some regulatory mechanisms to accelerate the velocity by the scaffold protein that anchors kinesin to the cargo vesicle as suggested from our previous analysis of the APP-transport vesicles (Chiba *et al*., 2014).

By fitting of the modified force-relation model (Figure 4D) to the experimental data {*χ*\*, *v*} (Figure 4D), a single value of the effective temperature (equation (8)) for 79 different endosomes was estimated to be 4200±200 K. This does not literally mean that the temperature of the cytoplasm is 4200 K. Although the exact physical meaning of the effective temperature is still controversial (Hayashi and Sasa, 2004), it is often observed *T*_eff_ > *T* in non-equilibrium systems (Cugliandolo, 2011). The most plausible interpretation would be that the fluctuation process(es) that dominantly determine the drag force are actively driven by using energy 14 times higher than the thermal energy, which might reflect the complex *in vivo* environment. Here, the free energy obtained by a single ATP hydrolysis is about 20 *k*_B_*T*, which would give a reference for the energy scale of active processes in living cells.

### Fluctuation measurement for retrogradely transported endosomes in axons

The traces for the retrogradely transported endosomes were similarly analyzed. Traces with the segments of constant velocity that lasted for longer than 3 seconds were first examined to confirm the time constant for the convergence as ~ 50 ms, and then analyzed the remaining shorter traces which had segments of constant velocity that lasted for about 2 seconds (Figure 5A). The plots (119 runs) also clustered into several groups. The discrete distribution of the *χ* values at *Δt*=50 ms (the proxy for the convergent value *χ*\*) was statistically confirmed by the *k*-means clustering. Thus, the presence of FPU was also demonstrated with retrogradely transported endosomes. The number of FPUs showed a weak positive correlation to the endosome size (Figure 5B) as observed with the anterogradely transported endosomes (Figure 4B), which is consistent with the geometric constraints model (Figure 4B, the bottom panel) for the regulation of the number of FPUs on the endosome. Note that the discrete distribution of χ* also is additional experimental support that the effective temperature (equation (8)) does not vary much among these 119 vesicles from 112 neurons.

**Figure 5.**
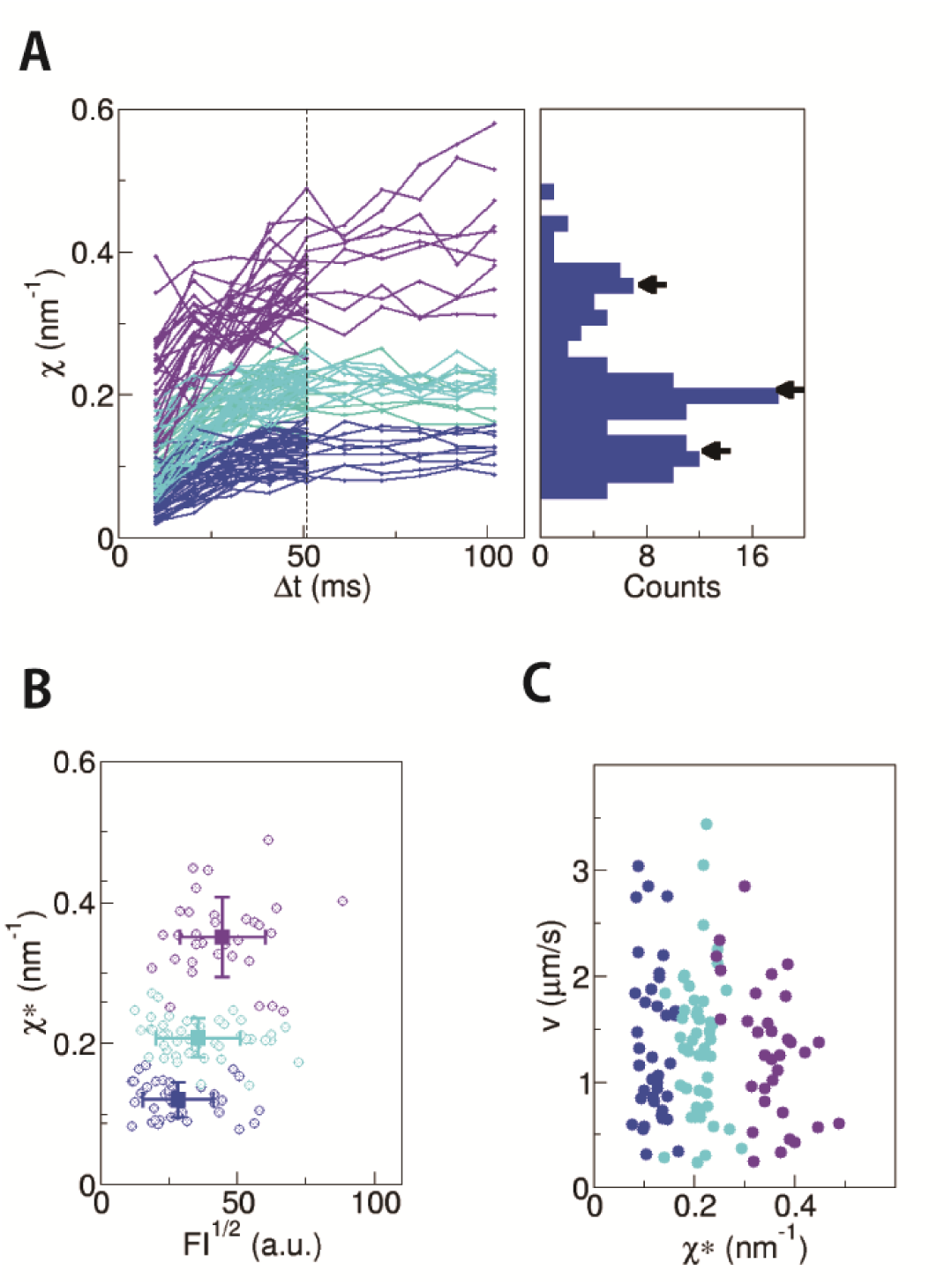
Measurement of χ for retrogradely transported endosomes in axon. **(A)** The traces of *χ* plotted against *Δt* for anterograde endosomes (n=119). They were classified into three clusters by *k*-means clustering (indicated with different colors. See Methods for the details). The distribution of *χ*\* is shown as a histogram. The positions of the cluster centers are indicated by arrows. **(B)** The fluctuation *χ*\* is plotted against the square root of fluorescence intensity (FI), which is a proxy for the radius of the endosome. The color of each data point is the same as in the left panels in (A), which reflects the number of FPUs. The average and the SD for each cluster are shown with square symbols with error bars. There was a weak correlation between *χ*\* and (FI)^1/2^ (*r*=0.33), as observed with anterograde endosomes (Figure 4B). **(C)** The *χ*\*-velocity relation. The colors for the data points are the same as the other panels.

The relation between the fluctuation χ* and the velocity for the retrograde vesicles (Figure 5C) showed similar distribution to the anterograde vesicles (Figure 4C), which would support that the same or similar phenomenological model for kinesin (Schnitzer *et al*., 2000) used in Figure 4C would be applicable to dynein as well. However, the force-velocity relations, especially the stall force for mammalian dynein *in vitro* are still controversial. If we assume the same or similar value of the effective temperature for the retrograde vesicles as the anterograde vesicles, the maximum force of a single dynein FPU would be around 10 pN, roughly same as a single kinesin. This maximum force value is apparently inconsistent with most *in vitro* studies that report only 1 pN for a single dynein molecule (Mallik *et al*., 2004; Hendricks *et al*., 2012; Rai *et al*., 2013). However, several groups independently reported that dynein can be activated to produce maximum force around 5 pN (Toba *et al*., 2006; Nicholas *et al*., 2015; Belyy *et al*., 2016), and there might be a mechanism to increase the maximum force to 10 pN in the cytoplasm. Thus, it would be difficult at this moment to examine the mechanochemistry parameters for dynein in the phenomenological model equations (M4-M6) and to compare them with the *in vitro* experiments.

## Discussion

In this study, we have demonstrated that the FT is practically useful for the non-invasive force measurement *in vivo*. The discrete distributions of χ* seen in Figure 4A and 5A along with the results in Figure 3 support that the proportional constant (*k*_B_*T*_eff_) between *F* and *χ*\* (equation (8)) does not vary much among ~200 vesicles analyzed here. The *χ*\*-*v* relation in Figure 4D fitted well to the force-velocity curve of kinesin in vitro by assuming a single value of *T*_eff_ (4200±200 K). Although the physical or mechanistic details that determine *T*_eff_ remain unclear, the results imply that the calibration for *T*_eff_ would not be necessary for each vesicle or neuron, but a single result of calibration can be applied to other vesicles. In other words, the degree of fluctuation *χ* can be used as a proxy for the force *F* in the case of cargo transport in neurons. The degree of fluctuation χ can be measured non-invasively even in living animals, and was shown to be very effective for the relative comparison of the force on the synaptic vesicles exerted by UNC-104 kinesin among mutant and wild type animals (Hayashi *et al*., 2018).

For the anterograde endosomes, the force value ~10 pN determined in this study is not only also consistent with the previous *in vivo* force measurements in macrophages (Hendricks *et al*., 2012), but also with the microrheology experiments (Wirtz, 2009). The effective viscosity *η*_eff_ in this study can be calculated from the force value and other parameter values (the diameter of endosome 2*r* =500 nm, and the velocity *v*=2 μm/s) to be about 1000 cP by using the relation *F* = 6*πη*_eff_*rv*. This estimate is 1000 times higher than water, but is consistent with the previous measurement in the cytoplasm with tracers of similar sizes to the endosomes (Wirtz, 2009). This 1000 times difference is explained that the effective viscosity includes the effect of the surrounding environments as well as the simple collisions with solvents, as discussed in the Theory section.

For the retrograde endosomes, the velocity distributions (Supplementary Figure S4), and the size distributions (the distributions of fluorescence intensities) (Figure 4B and 5B) were similar to the anterograde endosomes. The effective viscosity would reflect the environment of the axonal cytoplasm, which is common to both the anterograde and the retrograde endosomes. Then, the drag force exerted on the retrograde endosomes would be similar to the anterograde endosomes (Stokes’ law). Namely, the maximum force by a single FPU of dynein would not differ much from kinesin (around 10 pN). This force value is about 10 times higher than the previous *in vitro* studies (Mallik *et al*., 2004; Hendricks *et al*., 2012; Rai *et al*., 2013). Some mechanisms are expected to activate dynein in the cytoplasm. The average number of dynein molecules on a single vesicle (Hendricks *et al*., 2010) is two times larger than the average number of retrograde FPUs in this study. Two dimers of dynein might serve as a single FPU and produce force in a cooperative and collective manner (Torisawa *et al*., 2014). Indeed, some dynein adaptors such as BICDR and HOOK3 are recently reported to recruit two dynein dimers as a unit so that the unit can move at faster velocity and produce larger force (Urnavicius *et al*., 2018). Of course, another possibility not excluded is that only half of the dynein molecules on the endosome might be activated by some other mechanisms to produce force up to 10 pN.

In summary, we have established a FT-based method to estimate the drag force exerted on the vesicles transported in living cells by analyzing only their movement. Unlike other existing methods for force measurement, it is fully passive and non-invasive. We used vital staining with a fluorescent dye for a selective visualization of endosomes, but DIC or phase-contrast imaging of unstained samples can be used as well. Thus, this non-invasive method would serve as a powerful and versatile tool for basic research in the field of intracellular transport, as well as some potential applications for the examination of the molecular motor functions in clinical samples.

## Materials and Methods

### Reagents

All reagents were purchased from Wako or Sigma-Aldrich, unless otherwise stated.

### Primary culture of neurons

Superior cervical ganglions (SCGs) isolated from 3-week old ICR mice (male) were enzymatically treated in 0.5 % trypsin (Sigma) followed by 2 hr treatment with 0.5 % collagenase (Worthington). Dissociated cells were rinsed with DMEM/F12 containing 10 % heat inactivated bovine serum (Life Technologies), and plated onto Matrigel (BD-Biosciences)-coated glass-bottom dish (Matsunami). The neurons were cultured for two to four days with a DMEM/F12 medium supplemented with 10 % heat inactivated bovine serum and 200 ng/ml 2.5S nerve growth factor. All the animal experiments were conducted in compliance with the protocol which was approved by Institutional Animal Care and Use Committee, Tohoku University.

### Observation of endosomes and image analysis

The neurons were stained for 10 min with 100 nM DiI (DiIC18(3): 1,1’-Dioctadecyl-3,3,3’,3’-tetramethylindocarbocyanine perchlorate, Life Technologies), and then observed with a fluorescent microscope (IX71, Olympus) equipped with a heating plate (CU-201, Live Cell Instrument). The images of the motile endosomes were obtained with a 100x objective lens (UPlanFL 100x/1.3, Olympus) and an EMCCD camera LucaS (Andor) at 98 frames per second at 37 °C. The center position of each endosome was determined from the recorded image using ImageJ (Rasband, 1997), and the displacement from the position in the first frame was calculated for each frame. Here we focused on the displacement along the direction of the motion *X*(*t*). The data were collected from 34 preparations (culture dishes). 79 endosomes from 76 different cells for anterograde, and 119 endosomes from 112 different cells for retrograde were investigated. The cells for observation were chosen randomly after visual inspection, and the trajectories with longer than 2 second constant velocity run(s) were selected for the analyses. The constant velocity segment was selected by fitting the trajectory with a constant velocity movement, so that the residual does not exceed the variance perpendicular to the movement. The accuracy of the position measurement was verified with fluorescent beads with a similar size and fluorescent intensity to the endosomes (300 nm latex bead, Polyscience). The standard deviation of the position of the bead tightly attached to the glass surface was 8.4±0.4 nm (5 different beads in 2 independent preparations), which is much smaller than the displacement between frames *ΔX* (=*X*(*t*+*Δt*)-*X*(*t*)) analyzed in this study and would affect the accuracy of the fluctuation measurement by less than 10% CV, within the range of the estimation errors in the fluctuation (Supplemental Figure S1).

### Preparation of permeabilized and reactivated neurons

For some experiments, the plasma membrane of the neuron was permeabilized to control the cytoplasmic ATP concentration (Okada *et al*., 1995). Clarified brain homogenate was used to compensate for the loss of cytoplasmic components after membrane permeabilization. Mouse brain was cleaned in ice-cold PBS and homogenized with three volumes of KHMgE buffer (K-acetate 115 mM, Hepes 20 mM, MgCl2 1 mM, EGTA 1 mM, pH 7.4) supplemented with protease inhibitor cocktail (Complete EDTA-free, Roche). The homogenate was clarified by the successive centrifugation at 1,000 xg 10min and 100,000 xg 1 hr. The brain cytosol thus prepared was aliquoted and snap frozen with liquid nitrogen. The assay buffer was prepared just before use by mixing the brain cytosol with equal volume of KHMgE buffer supplemented with ATP regeneration system (0.125 mM or 0.0125 mM ATP, 10 mM creatine phosphate, 8 U/ml creatine phosphokinase), protease inhibitor cocktail (Complete EDTA-free) and 5 mM beta-mercapto ethanol. The neurons were first rinsed with KHMgE, then followed by 8 min incubation with the assay buffer containing 0.01 mg/ml digitonin. Fluorescent dextran (VECTOR) was used to examine the membrane permeabilization after digitonin treatment. The data were collected from 11 preparations (culture dishes) for [ATP]=125 μM (62 endosomes from 53 cells) and 6 preparations (culture dishes) for [ATP]=12.5 μM (30 endosomes from 23 cells).

### Analysis of fluctuation using FT

The value of *χ* is defined as

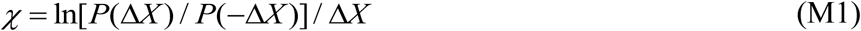

from the distribution, *P*(*ΔX*), of the displacement *ΔX* =*X*(*t*+*Δt*)-*X*(*t*). Since the noise was confirmed to be Gaussian (Figure 2C, D), *P*(*ΔX*) was fitted with a Gaussian function

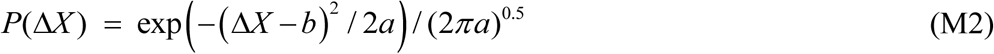

where the fitting parameters *a* and *b* correspond to the variance and the mean of the distribution. By substituting equation (M2) to equation (M1),

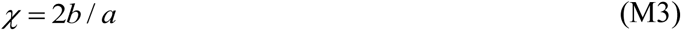

Thus, χ was calculated as 2*b*/*a* for each *P*(*ΔX*) for various interval *Δt* from 10 ms to 100 ms. The values for *a* and *b* were also estimated from the cumulative Gaussian distribution of *ΔX*, and directly as the sample variance (*a*=<(*ΔX*-<*ΔX*>)^2^>) and the average (*b*=<*ΔX*>). These two estimations provided the same values of *a* and *b* within the error of *χ* (Supplemental Figure S1). The converged value *χ*\* was determined by plotting χ against *Δt* as shown in Figure 2E (*χ*\* =*χ* at 51 ms in Figure 4, 5)

### *k*-means clustering

The *χ*-*Δt* plots for the anterograde and retrograde endosomes (Figure 4A, 5A) were classified statistically by using a *k*-means clustering method using a program package R with a library “cluster”. First, the number of clusters *k* was determined by calculating AIC (Akaike’s Information Criterion) for *χ*-*Δt* plots. In the case of the anterograde *χ*-*Δt* plots (Figure 4A), AIC values were ‐122.1 for *k*=2, ‐134.6 for *k*=3, ‐140.2 for *k*=4, and ‐134.4 for *k*=5. In the case of the retrograde *χ-Δt* plots (Figure 5A), AIC values were ‐232.3 for *k*=2, ‐246.6 for *k*=3, ‐247.0 for *k*=4 and ‐245.3 for *k*=5. From these AIC values along with the Gap statistics, the most probable values of *k* were determined as *k* =4 for anterograde and *k*=3 for retrograde, respectively. The initial value for the *k*-th cluster center trajectory χ_c_^k^ was chosen as the *k*-th peak value of *χ*\* (the arrows in Figure 4A, 5A). Each trajectory of *χ* was classified to the cluster based on the mean square deviation from χ_c_^k^. Then χ_c_^k^ was renewed as the mean of the trajectories classified to that cluster. This procedure was repeated until convergence.

### Analysis of the force-velocity relation

We adopted the widely-accepted model of kinesin-1 (Schnitzer *et al*., 2000) for the analysis of the measured force-velocity relations. In this model, velocity is expressed as the function of ATP concentration as

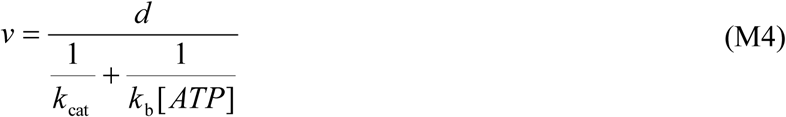

where *d* is the step size (=8 nm). *k*_cat_ and *k*_b_ are the catalytic turnover rate and the apparent second order rate constant for ATP binding (=ratio of *k*_cat_ and Michaelis-Menten constant *K*_M_). The load (*F*) dependencies are introduced as

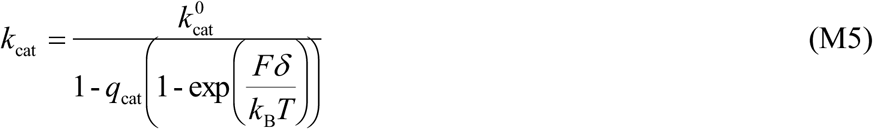

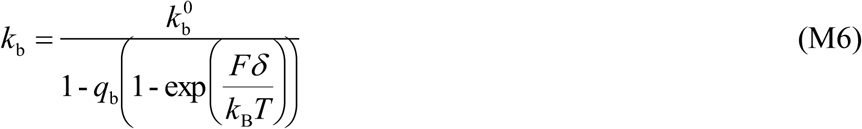

The values for the parameters *k*_cat_ and *k*_b_ were taken from the reported *in vitro* results (Schnitzer *et al*., 2000): *q*_cat_=0.0062, *q*_b_=0.04 and *δ*=3.7 nm.

The ATP concentration in the cytoplasm is around 4 mM. Since it is much higher than *K*_M_ which is around 50 μM for both kinesin and dynein, the inaccuracy in the ATP concentration does not affect the results. *k*_cat_^0^ was determined from the maximum velocity of endosome observed in our experiments (3.9 μm/s for anterograde). Namely, *k*_cat_^0^ = *v*_max_/*d* (488/s).

From the fitting of equations (M4-M6) to the experimental data {*χ*\*, *v*} in Figure 4D, the proportional constant (equation (8)) between *F* and *χ*\*

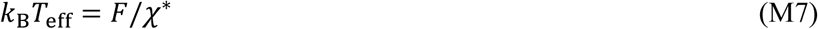

was estimated to be 58±3 pN nm (=14 *k*_B_*T*). Namely, the effective temperature *T*_eff_ was estimated to be 4200±200 K.

### Purification of kinesin for in vitro assay

As described previously (Okada and Hirokawa, 1999), a constitutive active dimer construct of mouse kinesin-1 (KIF5C 1-560aa) was subcloned into a plasmid vector pET21B (EMD biosciences). An *in vivo* biotinylation tag, BCCP (biotin carbonyl carrier protein, Promega) was inserted to the C-terminus of the construct. The plasmid was introduced into bacterial cell BL21(DE3)RIL (Agilent). The transformant was cultured with 2x YT medium supplemented with 30 mM phosphate buffer (pH 7.4) at 37 °C to mid-log phase (OD600=1.0). The culture was cooled down to 23 °C, and the protein expression was induced by adding 0.1 mM (final concentration) isopropyl β–D-1-thiogalactopyranoside (IPTG). The bacterial cells were collected 5 hrs after induction, and rinsed with ice-cold phosphate-buffered saline (PBS) supplemented with phenylmethyl-sulfonyl fluoride (PMSF).

Then, the bacterial cells were resuspended with five volumes of buffer A (HEPES 50 mM, potassium acetate 500 mM, magnesium acetate 5 mM, imidazole 10 mM, pH 7.4 adjusted with KOH) supplemented with ATP 0.1 mM and the following protease inhibitors: Pefabloc SC 1 mM, Leupeptin 20 μM, Pepstatin A 10 μM, N±-p-tosyl-L-arginine methyl ester (TAME) 1 mM. The bacterial cell wall was solubilized with lysozyme (2 mg/ml). DNase I (10 μg/ml) was added to reduce viscosity by the bacterial genomic DNA. Then bacterial cells were broken by sonication.

The soluble protein was recovered by centrifugation at 20,000 xg for 30 min, and was applied to the immobilized metal affinity chromatography column TALON (Takara). The protein was eluted with buffer B (PIPES 20 mM, imidazole 250 mM, magnesium sulfate 2 mM, EGTA 1 mM) supplemented with ATP 0.1 mM and protease inhibitors. The peak fractions were pooled and stored at ‐80°C after snap-freezing in liquid nitrogen.

### Preparation of microtubules

Tubulin was purified by the high-molarity PIPES buffer method (Castoldi and Popov, 2003) with modifications (Yajima *et al*., 2012). Porcine brains were cleaned by removing meninges, blood clots and vessels in washing buffer (PIPES 50 mM, PMSF 5 mM, pH 6.8). They were homogenized in a pre-chilled Waring blender with PEM buffer (PIPES 100 mM, EGTA 1 mM, MgCl_2_ 1 mM, pH 6.8 adjusted with KOH) supplemented with PMSF 0.5 mM, Leupeptin 2 µM and DTT 0.5 mM. After clarification with centrifugation at 15,200 xg, 60 min, microtubules were polymerized by warming the supernatant to 37 °C after supplementation with MgATP 1 mM, MgGTP 0.5 mM and glycerol. The polymerized microtubules were collected by ultracentrifugation at 100,000 xg 37 °C. Then, they were depolymerized in ice-cold P_1000_EM buffer (PIPES 1,000 mM, EGTA 1 mM, MgCl_2_ 1 mM, pH 6.8 adjusted with KOH) at 0 °C. The supernatant was collected by ultracentrifugation at 100,000 xg, 4 °C. The polymerization and depolymerization cycles were repeated four times, and the final supernatant was pooled and stored in liquid nitrogen.

TMR (tetramethyl rhodamine)-labeled microtubules were prepared as follows. Microtubules were polymerized in PEM buffer supplemented with 1 mM GTP at 37 °C. Then, 5-(and-6)-Carboxytetramethylrhodamine, succinimidyl ester (Life Technologies) was added at 5-10 molar excess. Labeled microtubules were separated from free dye by ultracentrifugation through glycerol cushion, and were resuspended with ice-cold PEM buffer. The microtubules were depolymerized by cooling down the solution to 0°C, and the supernatant was collected by ultracentrifugation at 100,000 xg, 4 °C. The labeling efficiency was measured spectroscopically, and the microtubules were stored in liquid nitrogen.

### Bead assay

For the bead assay, the carboxy-modified fluorescent 0.5 µm latex bead (Life Technologies) was biotinylated with (+)-biotinyl-3,6,9,-trioxaundecanediamine (Amine-PEG3-biotin, Pierce) using condensation agent DMT-MM(4-(4,6-dimethoxy-1,3,5-triazin-2-yl)-4-methyl-morpholinium). The purified recombinant kinesin dimer was immobilized on the bead surface via streptavidin (Sigma) in assay buffer (PIPES 80 mM, magnesium acetate 5 mM, EGTA 1 mM, ATP 2 mM, casein 0.5 mg/ml, taxol 10 µM, β-mercaptoethanol 10 mM, catalase 0.1 mg/ml, glucose 10 mM, glucose oxidase 0.05 mg/ml, pH 6.8). Diluted, TMR-labeled microtubules were absorbed to the glass surface of the flow cell chamber, and the remaining surface was coated with a biocompatible polymer Lipidure-BL-103 (NOF, Tokyo, Japan). Then, the kinesin-coated beads were injected into the chamber. The optical tweezers instrument is based on the inverted microscope IX2 (Olympus). The beam of near infra-red laser (1064 nm BL-106C, Spectra-Physics) was collimated to fill the back aperture of the objective lens (PlanApo 60x/1.40, Olympus). The bead trapped at the focus was illuminated with green laser (532 nm, 400 mW, Genesis CX, Coherent), and its image was projected to EMCCD camera iXon DU-860D-CS0-#BV (Andor). The stiffness of the trap was 0.1 pN/nm. The images were recorded at the speed of 400 frames per second at 22 °C. The constant velocity segments (n=45) used in the analysis (Figure S3) were cut from 31 runs from 5 different bead assays.

## Acknowledgments

We thank N. Sawairi and M. Tomishige for initial stages of experiments; S. Xu, J. Asada, M. Komeno, M. Kakiuchi and K. Ito for their technical and secretarial assistance; W. Kylius for editing the manuscript. This work was supported by AMED under grant number JP17gm5810009, by the Ministry of Education, Culture, Sports, Science and Technology, Grant-in-Aid for Scientific Research (KAKENHI) (grant numbers 26104501, 26115702, 26310204, 16H00819) to K.H, as well as the following grants to Y.O.: KAKENHI (grant numbers 24659092, 25113723, 25293046, 26115721, 26650069, 15H01334, 17K19511), the Uehara memorial Foundation, Takeda Science Foundation, and the Naito Foundation. Y.T. was supported by “Program for Leading Graduate Schools” of MEXT.

## Supplementary Materials

**Legends for Supplemental Figures S1-S5**

**Figure S1.**
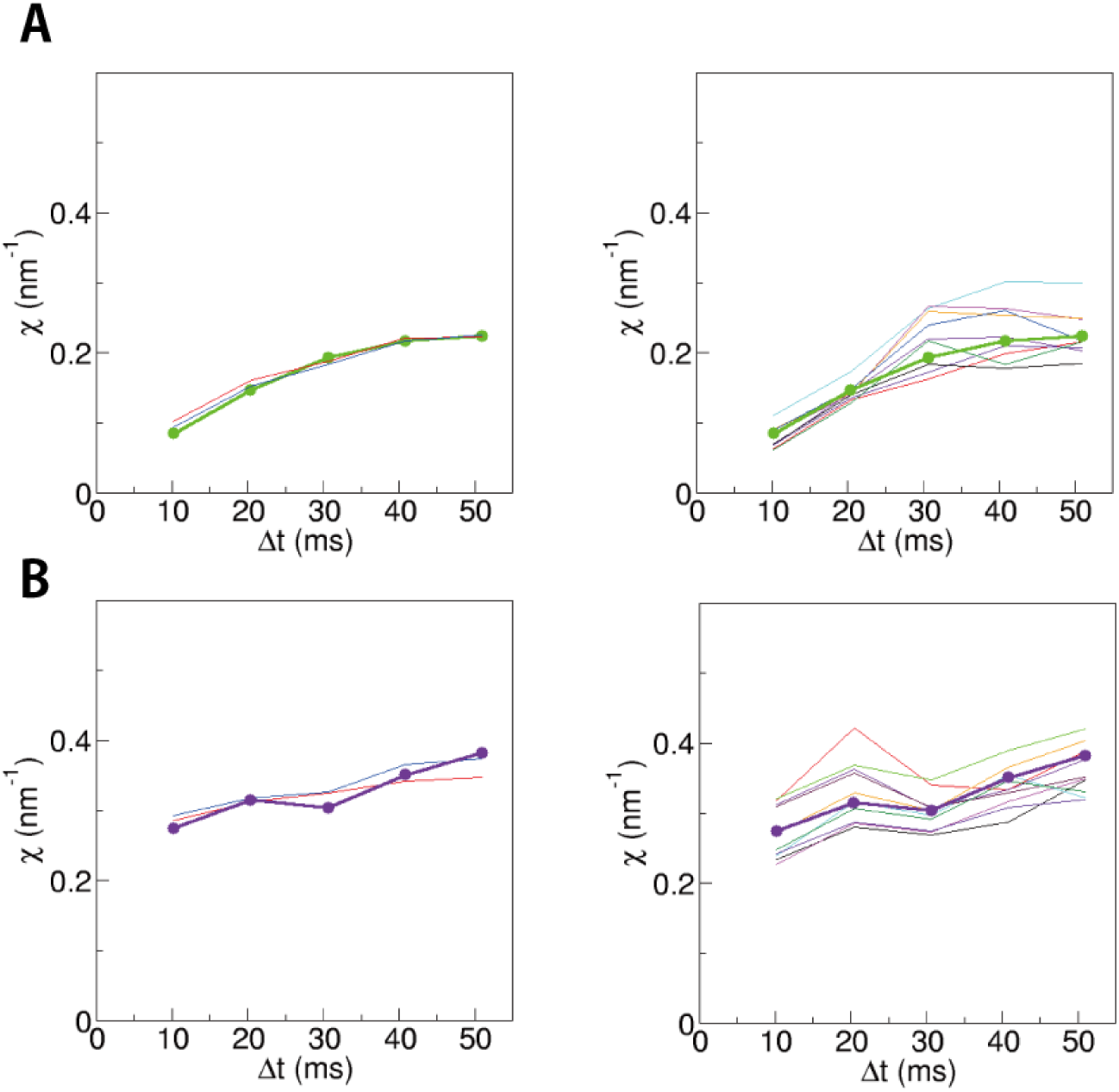
Evaluation of the errors in the estimation of the fluctuation χ. Panels (**A**) and (**B**) represent an example of anterograde and retrograde vesicles, respectively. The left panels show the comparison of the method to estimate the Gaussian distribution from the observed *ΔX* values. The thick curves represent the estimation from the histogram of *ΔX*, which can be biased by binning. The thin blue curves are based on the fitting of the cumulative Gaussian distribution. The thin red curves show the results by simply calculating the average and the variance of *ΔX*, which can be biased by the tails of the distribution. The right panels show 10 sample results of the bootstrapping confirmation for the errors in the estimation. These bootstrapping results gave an estimate for the errors in the estimation of the fluctuation *χ* as 10% CV, which is consistent with the expected errors from the accuracy of the position measurement (8 nm).

**Figure S2.**
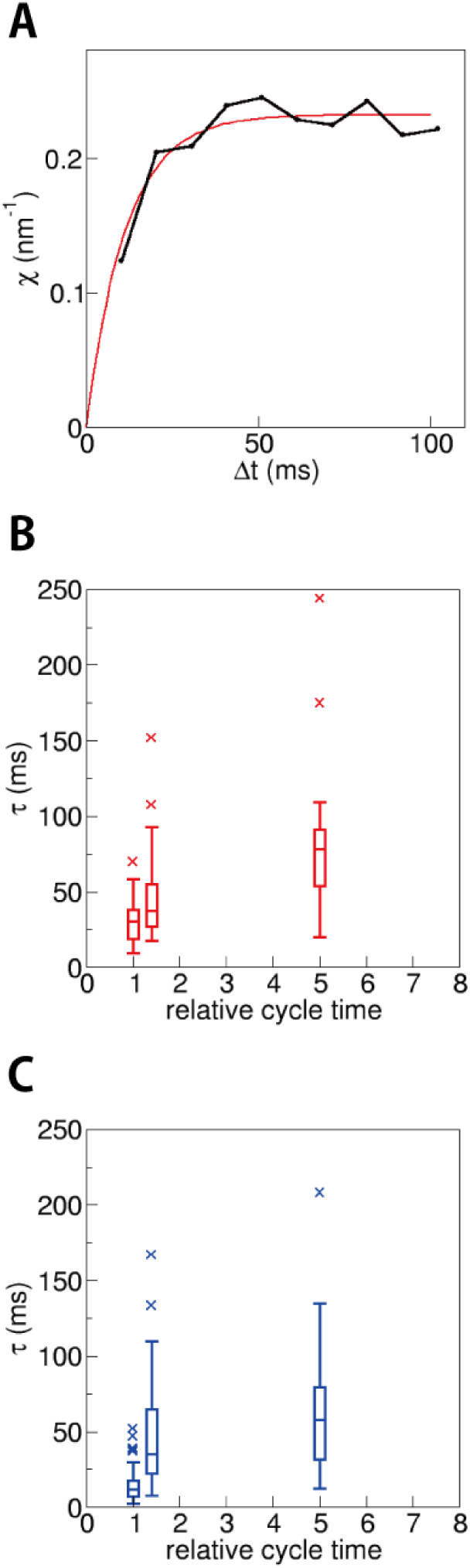
Analysis of the relaxation time. (A) The relaxation time *τ* was estimated by fitting of the function *c*(1-exp(*-Δt/τ*)) to *χ*, where *c* is a constant. In this paper, the converged value of *χ*\* was calculated as *χ* at *Δt*=51 ms, noting that *χ* at *Δt*=51 ms is the same as *c* within the error of *χ*. **(B, C)** The fluctuation of the endosomes was analyzed in permeabilized and reactivated neurons, in which the ATP concentration was maintained at 125 μM and 12.5 μM. Both anterograde (B) and retrograde (C) endosomes showed slower relaxation when ATPase cycle was slowed down. The relative cycle time was calculated as 1+*K*_m_/[ATP], where the Michaelis-Menten constant *K*_m_ was assumed to be 50 μM as an approximate value for both kinesin and dynein. The data are shown as box-and-whisker plots. 35 anterograde and 27 retrograde vesicles (from 29 and 24 cells) were analyzed for [ATP]=125 μM. 15 vesicles each (from 9 and 14 cells) for [ATP]=12.5μM.

**Figure S3.**
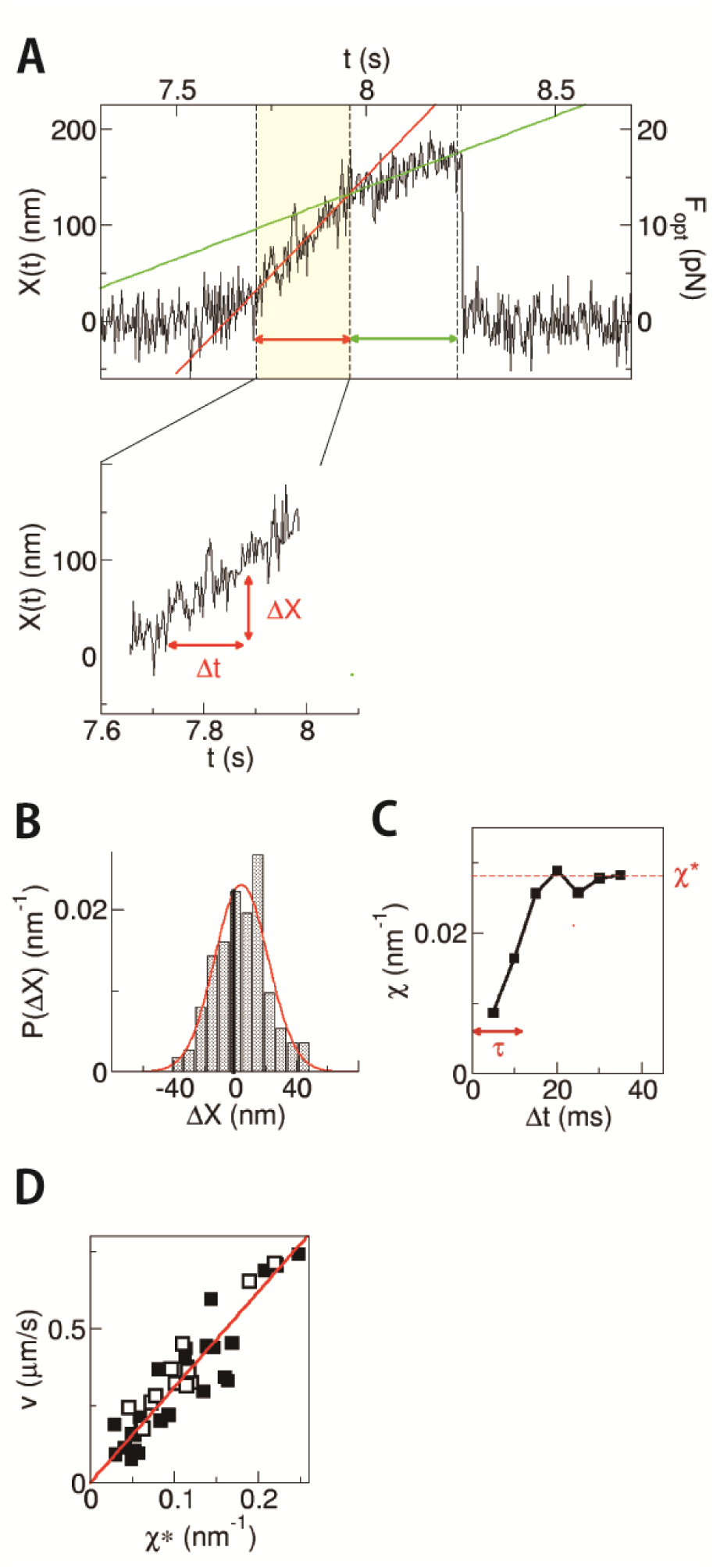
Relation between velocity and χ^*\**^ for the kinesin-coated bead. **(A)** An example of the trace of displacement *X*(*t*) of the bead, carried by kinesin motors, obtained in the *in vitro* experiment using an optical tweezers instrument (the stiffness of the trap was 0.1 pN/nm). Here loads on the beads were applied by optical tweezers(*F*_opt_) mimicking the high viscous effect acting on cargo vesicles in cells. Segments for the analysis in the graph indicated with red and green were determined by fitting the trajectory to the constant velocity movement. (inset) Analysis of fluctuation, calculated as *ΔX*=*X*(*t*+*Δt*)*-X*(*t*). **(B)** Distribution of *ΔX* at *Δt*=30 ms fitted by a Gaussian function. **(C)** Relaxation of *χ*. *χ*\* and *τ* denote the value of convergence and the time constant, respectively. **(D)** Linear relation between *χ*\* and *v* for the traces with the stall force <12 pN (closed squares (n=29)) and >12 pN (open squares (n=16)). The data points aligned linearly, supporting equation (9). The same proportional relation for the open and closed symbols indicates that the proportional coefficient (*k*_B_*T*_rff_/*Γ*) was not sensitive to the number of motors attached to the beads.

**Figure S4.**
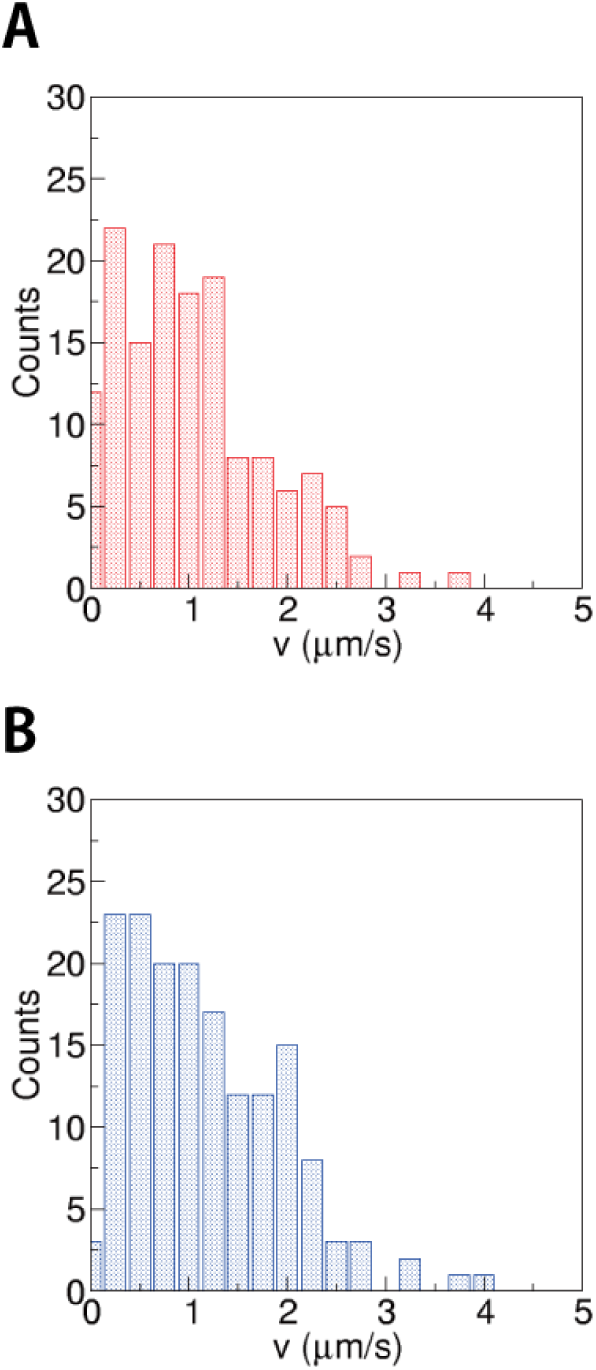
Velocity distributions of endosomes. Velocity distributions of endosomes for anterograde transport **(A)** and retrograde transport **(B)**. Only the segments of constant velocity movement with durations longer than 2 s were analyzed (e.g. rectangles depicted in Figure 3B). The mean velocities were 1.2 ± 0.7 μm/s (mean ± SD, n=145) for anterograde transport, and 1.3 ± 0.8 μm/s (n=163) for retrograde transport, respectively. It should be noted that the velocities distributed more continuously without clear distinct peaks, which appears very differently from the distributions of *χ*\* (Figure 4A, 5A).

